# The *Helicobacter pylori* biofilm involves a multi-gene stress-biased response including a structural role for flagella

**DOI:** 10.1101/412239

**Authors:** Skander Hathroubi, Julia Zerebinski, Karen M. Ottemann

## Abstract

*Helicobacter pylori* has an impressive ability to persist chronically in the human stomach. Similar characteristics are associated with biofilm formation in other bacteria. The *H. pylori* biofilm process, however, is poorly understood. To gain insight into this mode of growth, we carried out comparative transcriptomic analysis between *H. pylori* biofilm and planktonic cells, using the mouse colonizing strain SS1. Optimal biofilm formation was obtained with low serum and three-day growth, conditions which caused both biofilm and planktonic cells to be ∼80% coccoid. RNA-seq analysis found that 8.18% of genes were differentially expressed between biofilm and planktonic cell transcriptomes. Biofilm-downregulated genes included those involved in metabolism and translation, suggesting these cells have low metabolic activity. Biofilm-upregulated genes included those whose products were predicted to be at the cell envelope, involved in regulating a stress response, and surprisingly, genes related to formation of the flagellar apparatus. Scanning electron microscopy visualized flagella that appeared to be a component of the biofilm matrix, supported by the observation that an aflagellated mutant displayed a less robust biofilm with no apparent filaments. We observed flagella in the biofilm matrix of additional *H. pylori* strains, supporting that flagellar use is widespread. Our data thus supports a model in which *H. pylori* biofilm involves a multi-gene stress-biased response, and that flagella play an important role in *H. pylori* biofilm formation.

**IMPORTANCE:** Biofilms, communities of bacteria that are embedded in a hydrated matrix of extracellular polymeric substances, pose a substantial health risk and are key contributors to many chronic and recurrent infections. Chronicity and recalcitrant infections are also common features associated with the ulcer-causing human pathogen *H. pylori.* However, relatively little is known about the role of biofilms in *H. pylori* pathogenesis as well as the biofilm structure itself and the genes associated with this mode of growth. In the present study, we found that *H. pylori* biofilm cells highly expressed genes related to cell envelope, stress response and those encoding the flagellar apparatus. Flagellar filaments were seen in high abundance in the biofilm. Flagella are known to play a role in initial biofilm formation, but typically are downregulated after that state. *H. pylori* instead appears to have co-opted these structures for non-motility roles, including a role building a robust biofilm.

## INTRODUCTION

*H. pylori* has been co-evolving with humans for tens of thousands of years (1). During this time, it has adapted to survive the hostile environment of the stomach and evade the immune system, allowing it to persist for the life of the host (2). *H. pylori* colonizes gastric epithelial surfaces and within the thin layer of mucus near the cells (3). More recently, *H. pylori* was found to colonize within gastric glands, repeated invaginations of the gastrointestinal tract, which may provide the bacteria a favorable niche (4,5). Even though most infections are asymptomatic, *H. pylori* persistence is considered a major risk factor for gastric and duodenal ulcers, gastric Mucosa-Associated Lymphoid Tissue (MALT) lymphoma, and gastric adenocarcinoma (6). *H. pylori* infections remain difficult to treat, and when left untreated (7), 1-2% progress to gastric cancer (8,9).

*H. pylori* possesses several mechanisms to escape the challenging environment of the stomach where the pH is around 2. These include urease production, flagellar motility, and chemotaxis, which are all required for the initial and sustained colonization of the gastric epithelial surface (10). Urease catalyzes the hydrolysis of urea, which is abundant in stomach, into bicarbonate and ammonia and thus raises the pH to near neutral (10). pH elevation decreases the viscoelastic properties of mucus gel and improves the motility of *H. pylori*, which can then swim away from the lumen to reach safer niches including those close to the gastric epithelial surface (11). *H. pylori* forms microcolonies at the cell surface *in vitro* (12,13) as well as within gastric glands (5). This microcolony mode of growth may be consistent with the bacteria being in a biofilm-growth mode.

Biofilms are dense aggregates of microorganisms attached to a surface and embedded in an extracellular polymeric matrix (14). In contrast with the other mode of bacterial growth, free-floating or planktonic, biofilm cells tend to be more tolerant towards antimicrobials and host immune responses (14,15). Biofilms are also frequently associated with chronic disease including pneumonia in cystic fibrosis patients, Lyme disease, and chronic otitis media (16-18). In those chronic diseases, biofilm growth is considered to be a survival strategy used by pathogens to escape antimicrobial therapies, avoid clearance by the immune system, and to persist for the lifetime of the host.

Chronicity and recalcitrant infections are also common features associated with *H. pylori* (19). Yet, the role of biofilm growth in promoting *H. pylori* persistence is still not clear (20). The first suggestion of biofilm formation by *H. pylori* during colonization of the human gastric mucosa was found using biopsies and scanning electron microscopy (SEM) analysis (20-22). These studies demonstrated that gastric biopsy samples from *H. pylori*-positive patients showed dense layers of bacteria aggregated and attached to the mucosal surface. The bacteria were consistent in appearance to *H. pylori* with cells in both the spiral and coccoid morphologies. The same bacterial-appearing structures were absent in *H. pylori-*negative patients, however there has not yet been conclusive evidence showing that *H. pylori* forms a biofilm *in vivo*.

*H. pylori* has been well documented to form a biofilm *in vitro*. The first report of *in vitro* biofilm formation by *H. pylori* was described to occur in clinical, laboratory, and mouse-adapted strains, and was observed at the air-liquid interface on glass coverslips when the bacteria were grown in Brucella broth (BB) supplemented with slightly lower than normal fetal bovine serum (FBS) (23). The biofilms were mainly composed of coccoid bacteria, with a minority of spiral and rod shaped ones (23). In subsequent reports, scientists analyzed the extracellular polymeric substance (EPS) of *H. pylori* biofilms and found proteomannans, LPS-related structures, extracellular DNA, proteins, and outer membrane vesicles (24,25).

Additionally, biofilm cells have been shown to exhibit high resistance *in vitro* to clarithromycin, which is one of the common antibiotics used to treat *H. pylori* infection (26). The minimum inhibitory concentration (MIC) and the minimum bactericidal concentration (MBC) were increased by 16-and 4-fold, respectively, in the biofilm cells as compared to planktonic ones (26). However, despite the growing evidences of *H. pylori* biofilm formation both *in vitro* and *in vivo* (20-22,24), little is known about the genes involved in biofilm formation. We thus sought to characterize *H. pylori* biofilm and investigate global transcriptional changes during biofilm formation, with a particular focus on *H. pylori* strain SS1 because it is able to colonize mice and thus will be able to serve as a model for biofilm formation *in vivo*.

## RESULTS

### Biofilm formation and growth condition

*H. pylori* strain SS1 has been extensively used as a murine model of *H. pylori* infection. *H. pylori* SS1 biofilms, however, are difficult to detect when the bacteria are grown in standard nutrient-rich media routinely used for *H. pylori* culture. A previous study reported that *H. pylori* biofilm formation was significantly dependent on the growth media used (27). We thus evaluated the ability of the *H. pylori* SS1 strain to form a biofilm using the crystal violet biofilm assay, and bacteria grown under varying growth conditions that included different growth media, incubation times, and concentrations of serum.

Using Brucella Broth (BB) media supplemented with 10% FBS (BB10), the condition usually used for *H. pylori* liquid growth, no biofilm was detected. We therefore explored lower amounts of serum, as these have been reported elsewhere to promote adhesion of *H. pylori* strain 26695 (28) and biofilm formation of reference *H. pylori* strain ATCC 43629 and clinical strains *H. pylori* 9/10 (27). While only a slight biofilm was observed when *H. pylori* SS1 strain was grown with BB supplemented with 6% FBS, a pronounced biofilm (*p*< 0.01) was detected in BB supplemented with 2% FBS (BB2) (Fig. 1A). The *H. pylori* growth rate was slightly reduced in BB2 compared with BB10, which suggests that the increase of biofilm formation was not due to increased growth (data not shown). HAMs F12 similarly only supported biofilm formation with low FBS percentages (Fig. 1B). Further experiments identified that three days of growth in BB2 led to the greatest amount of biofilm (Fig. 1C). These results thus suggest that BB media supplemented with 2% serum and growth for three days is an optimal condition for studying *H. pylori* SS1 biofilm formation.

**FIG 1.**
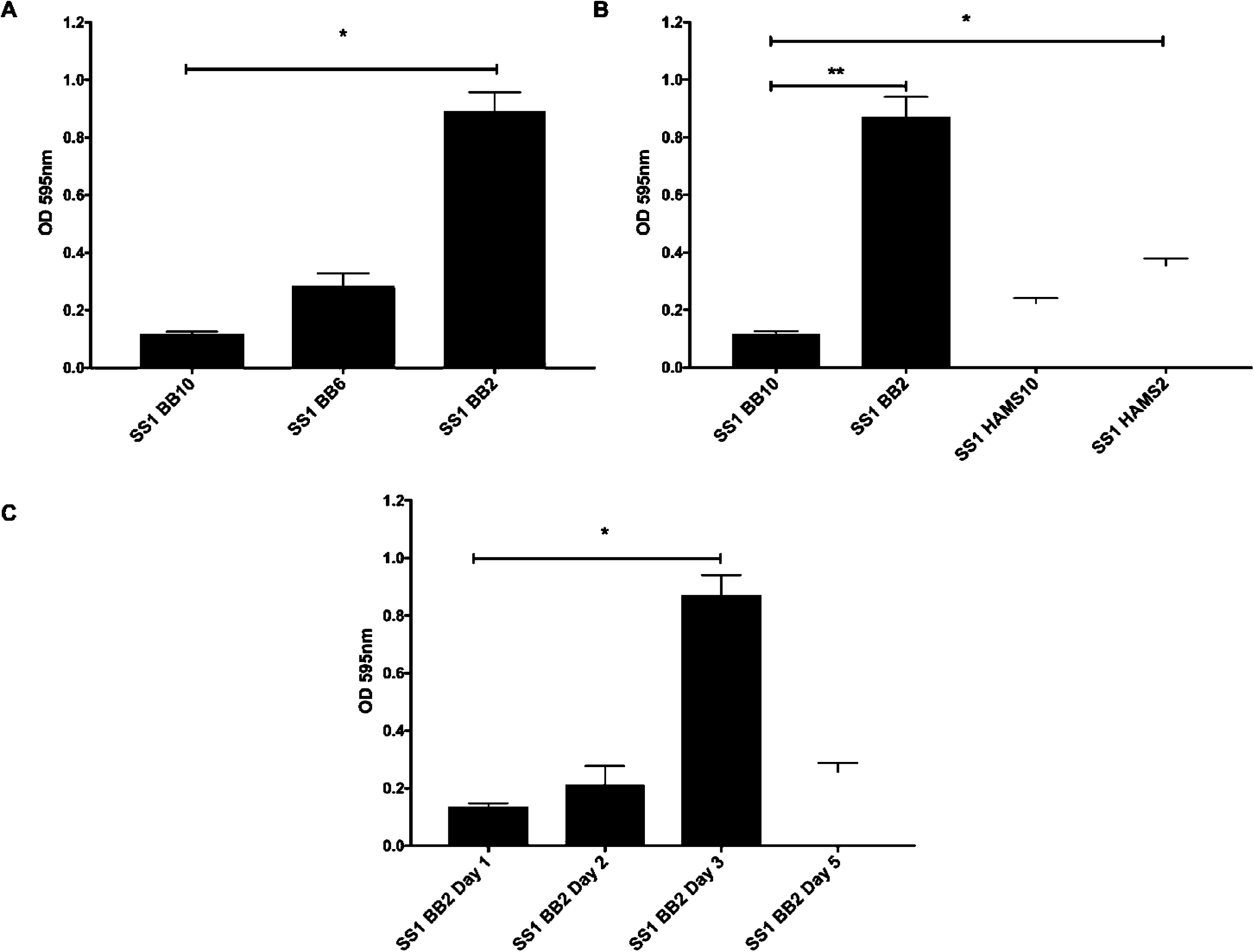
*H. pylori* SS1 forms robust biofilms after 3 days of growth in BB2. *H. pylori* strain SS1 was grown in the indicated media and biofilm formation was assessed by crystal violet absorbance at 595nm. **(A)** *H. pylori* SS1 was grown for 3 days in BB media supplemented with different concentration of FBS (BB10: 10%, BB6: 6% and BB2: 2%). **(B)** *H. pylori* SS1 was grown for 3 days in BB media or HAMS F12 supplemented with 10% or 2% of FBS. **(C)** *H. pylori* SS1 was grown in BB media supplemented with 2% FBS and biofilm formation was evaluated at different time points. Experiments were performed three independent times with at least 6 technical replicates for each. Statistical analysis was performed using ANOVA (*, *P* < 0.05 and **, *P* < 0.01).

### Biofilm characterization

To confirm and extend the results obtained with the crystal violet biofilm assay, biofilms of *H. pylori* SS1 were visualized by confocal laser scanning microscopy (CLSM) and staining with FilmTracer™ FM^®^1–43, a dye that fluoresces once inserted into the cell membrane. After three days of growth in BB2, we observed a thick bacterial biomass that non-homogeneously covered the surface, consistent with a well-developed biofilm (Fig. 2). Using z-stack images, the thickness of the *H. pylori* SS1 biofilm was determined to be 11.64 ± 2.63 μm^3^/μm (Supplementary Movie 1). As expected, *H. pylori* SS1 grown in BB10 did not form a biofilm that could be visualized with CLSM (data not shown).

To further characterize the EPS that composed the SS1 biofilm matrix, BOBO-3 and FilmTracer SYPRO Ruby biofilm matrix stains were used to stain extracellular DNA (eDNA) and extracellular proteins, respectively as described previously (29,30). Both of these molecules extensively stained the biofilm EPS, consistent with the idea that the *H. pylori* SS1 biofilm matrix contains significant amount of eDNA and extracellular proteins (Fig. 2B and C). Because these same molecules have been detected in other *H. pylori* strains, these results suggest that the *H. pylori* EPS is typically composed of eDNA and proteins (24,31).

**Fig 2.**
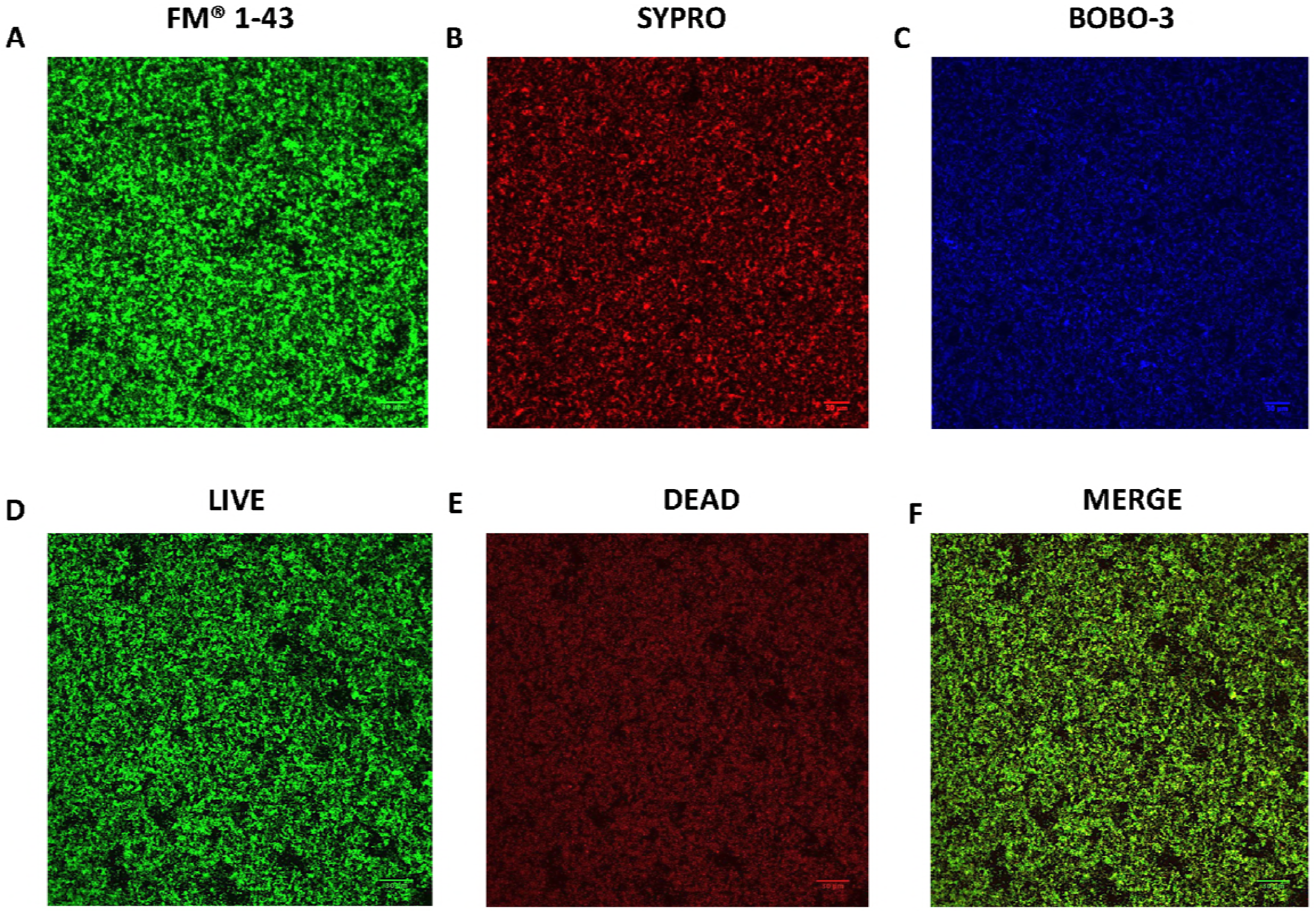
Confocal scanning laser microscopy (CSLM) images of *H. pylori* SS1 biofilm. Shown are representative CSLM images of 3 day-old SS1 biofilms grown in BB2 and stained with **(A)** FM 1-43 to stain total bacterial cells; **(B)** SYPRO RUBY to stain extracellular proteins, **(C)** BOBO-3 to stain extracellular DNA and **(D-F)** Live-Dead staining with live cells represented by the green-fluorescent SYTO 9, and dead/damaged cells by the red-fluorescent propidium. Scale bar = 30 µm

We also performed live-dead staining with the FilmTracer LIVE/DEAD biofilm viability kit, to define whether the biofilm cells were alive or dead. This approach revealed a subpopulation of dead or damaged cells, stained red, that appear to be homogeneously distributed within the live biofilm cells, which stained green (Fig. 2D-F). This result suggests that the *H. pylori* biofilm contains both live and dead cells.

To determine the importance of extracellular proteins and eDNA in the biofilm matrix of *H. pylori* SS1, we employed enzymatic treatment using DNAse I and proteinase K. Proteinase K treatment significantly dispersed pre-formed biofilms (*P* < 0.01) (Fig. 3). *H. pylori* pre-formed biofilms were, however, resistant to DNase treatments. These data suggest that DNA may play only a minor role in the biofilm matrix, however, extracellular proteins likely play an important role in the biofilm architecture of *H. pylori*, as has been reported in other *H. pylori* strains (24,31). These results suggest that many *H. pylori* strains, including SS1, use a protein-based biofilm matrix.

**Fig 3.**
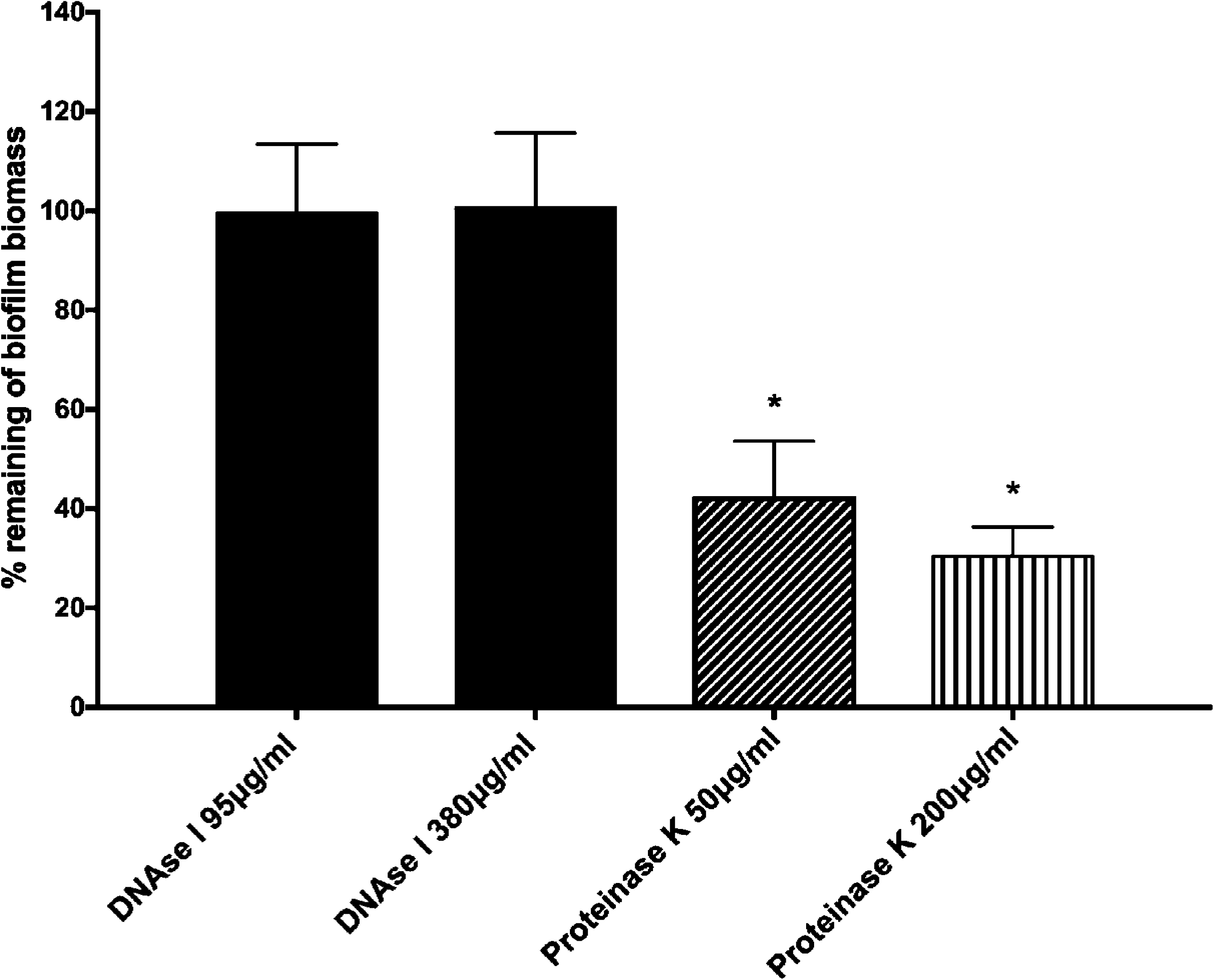
Effect of enzymatic treatments on pre-formed biofilms. *H. pylori* SS1 was allowed to form biofilms for three days in BB2. The media was then removed and replaced with either fresh media or media containing DNase I or proteinase K. Cells were re-incubated for 24 hours, and then analyzed for remaining biofilm using the crystal violet assay. Data shown here represent the percentage of remaining of biofilm compared to the untreated control. Experiments were performed three times independently with at least 8 technical replicates for each. Statistical analysis was performed using ANOVA *, *P* < 0.01 compared to the untreated control.

### Transcriptomic profiling of biofilm versus planktonic cells

To gain insight into the genes involved in *H. pylori* biofilm growth, we performed a transcript profiling experiment using RNA-seq. For this experiment, we grew *H. pylori* SS1 in BB2 in six well plates for three-days, and collected the free-floating planktonic cells and the bottom-attached biofilm ones from the same wells. We collected RNA from three biological replicates grown on two separate days. A total of 10-20 million reads per sample was generated by RNA-seq. These reads were then mapped to *H. pylori* SS1 complete reference genome (32), and revealed a clear clustering of the biofilm-grown cells in a distinct population compared to the planktonic ones (Fig. 4A). This transcriptomic analysis showed that 122 of 1491 genes (8.18%) were significantly differentially expressed (*p*< 0.01 and log2-fold change >1 or <-1) between *H. pylori* biofilm and planktonic populations (Fig. 4B and Fig. 5). 61 genes were significantly upregulated in biofilm cells compared to their planktonic counterparts, while another 61 were significantly upregulated in planktonic cells (Table 2 and 3). To validate the results obtained by this RNA-seq, the relative abundance of selected RNA transcripts was quantified by quantitative RT-PCR (qRT-PCR). Using this approach, we detected the same gene expression trend between qRT-PCR and RNA-seq, thus validating our results (Fig. 6). Below we discuss the most prominent of these genes, and what they suggest about the *H. pylori* biofilm growth state.

Our data suggest that biofilm cells may be less metabolically active than planktonic cells, based on the decreased expression of several genes involved in translation and ribosomal structure (Fig. 5, Table 3). Similarly, genes involved in metabolism, biosynthesis of cofactors, and urease were also down-regulated (Fig. 5, Table 3).

**Table 1.** Strains used in this study.

| *H. pylori* strain | KO Strain number | Description/genotype | Reference/source |
| --- | --- | --- | --- |
| SS1 WT |  | Wild-type strain | (62)/ J. O'Rourke |
| SS1 $\Delta fliM$ | KO1064 | $\Delta fliM::cat$ | This study (allele published in (41)) |
| SS1 $\Delta motB$ | KO536 | $\Delta motB2$ | (65) |
| G27 WT |  | Wild-type strain | (66)/ N. Salama, Fred Hutchison Cancer Research Center, Seattle |
| G27 *motB* | KO493 | *motB::aphA3* | This study |
| G27 *flgS* | KO688 | *flgS::TnCat* | (67) |
| G27 *fliA* | KO689 | *fliA::TnCat* | (67) |

**Table 2.** Up-regulated gene in *H. pylori* SS1 biofilm (cutoff ratio ≥ 1 log2 fold change and p-value <0.05) using RNA-seq analysis, grouped by functional role categories. Fold change represents the difference in gene expression between biofilm (n =3) and planktonic (n =3) populations.

| Locus Name | Putative identification | Fold Change |
| --- | --- | --- |
| **Cell envelope** |  |  |
| *flgL* (HPYLSS1_00284) | Flagellar hook-associated protein 3 | 7.64 |
| *flgK* (HPYLSS1_01062) | flagellar hook-associated protein 1 | 6.16 |
| *flgM* (HPYLSS1_01066) | Anti-sigma-28 factor | 3.94 |
| *flaG* (HPYLSS1_00586) | Polar flagellin G | 3.93 |
| *flaB* (HPYLSS1_00110) | Flagellin B | 3.52 |
| *flgE1* (HPYLSS1_00464) | Flagellar hook protein 1 | 3.07 |
| *flgB* (HPYLSS1_01503) | Flagellar basal body rod protein | 2.37 |
| *fliL* (HPYLSS1_00526) | Flagellar protein of unknown function | 2.25 |
| *fliK* (HPYLSS1_00653) | Flagellar hook-length control protein | 2.14 |
| *fliD* (HPYLSS1_00585) | Flagellar hook-associated protein 2 | 2 |
| *lpxB* (HPYLSS1_00467) | Lipid-A-disaccharide synthase | 3.06 |
| *lptB* (HPYLSS1_00622) | Lipopolysaccharide export system ATP-binding | 2.54 |
| *mltD* (HPYLSS1_01517) | Membrane-bound lytic murein transglycosylase D precursor | 2 |
| *pgdA* (HPYLSS1_00299) | Peptidoglycan deacetylase | 2.1 |
| HPYLSS1_00450 | Membrane protein | 3.78 |
| HPYLSS1_01378 | Outer membrane protein *homD* | 3.26 |
| HPYLSS1_01113 | Putative outer membrane protein | 2.91 |
| HPYLSS1_01021 | Outer membrane protein *homC* | 2.69 |
| HPYLSS1_01469 | Putative outer membrane protein | 2.52 |
| **Cellular process** |  |  |
| *cagE* (HPYLSS1_00705) | Type IV secretion system protein virB4/DNA transfer | 2.46 |
| *cagW* (HPYLSS1_00718) | CAG pathogenicity island protein CagW (cag10) | 2.31 |
| *cagL* (HPYLSS1_00710) | CAG pathogenicity island protein CagL (cag18) | 2.02 |
| *recR* (HPYLSS1_00636) | Recombination protein RecR | 3.7 |
| **HPYLSS1_00410** | DNA polymerase I | 2.08 |
| **HPYLSS1_01332** | CMP-N-acetylneuraminate-beta-galactosamide- | 2.2 |
| **Regulatory functions** |  |  |
| *hrcA* (HPYLSS1_00106) | Heat-inducible transcription repressor HrcA | 4.84 |
| *hspR* (HPYLSS1_00407) | Putative heat shock protein HspR | 2.09 |
| *crdR* (HPYLSS1_01312) | Two component response regulator CrdR | 2.03 |
| *rsfS* (HPYLSS1_01340) | Ribosomal silencing factor S | 2.16 |
| **Translation** |  |  |
| *ansA* (HPYLSS1_00615) | Putative L-asparaginase | 2.11 |
| *cbpA* (HPYLSS1_00408) | Curved DNA-binding protein | 2.19 |
| **HPYLSS1_00252** | Chaperone protein ClpB | 2.7 |
| **HPYLSS1_01332** | CMP-N-acetylneuraminate-beta-galactosamide | 2.2 |
| **Amino acid biosynthesis** |  |  |
| *porC* (HPYLSS1_01050) | Pyruvate synthase subunit PorC | 2.02 |
| **Fatty acid and phospholipid metabolism** |  |  |
| *acpS* (HPYLSS1_00527) | Holo-[acyl-carrier-protein] synthase | 2.89 |
| *fenF* (HPYLSS1_00085) | Malonyl CoA-acyl carrier protein transacylase | 2.26 |
| **Biosynthetic of cofactors, prosthetic groups and carriers** |  |  |
| *thiE* (HPYLSS1_00492) | Thiamine-phosphate synthase | 4.13 |
| *salL* (HPYLSS1_00914) | Adenosyl-chloride synthase | 2.87 |
| **DNA restriction, modification, recombination, and repair** |  |  |
| **HPYLSS1_00696** | Restriction endonuclease | 2.19 |
| **Transport and binding proteins** |  |  |
| ***metI* (HPYLSS1_01522)** | D-methionine transport system permease protein | 2.18 |
| **HPYLSS1_00805** | Putative ABC transporter ATP-binding protein | 2.02 |
| **Energy metabolism** |  |  |
| ***ansA* (HPYLSS1_00615)** | Putative L-asparaginase | 2.11 |
| **HPYLSS1_00772** | Pyrroloquinoline quinone biosynthesis protein | 7.5 |
| **Hypothetical proteins** |  |  |
| **HPYLSS1_00605** | hypothetical protein/Putative GTPase dynamin | 17.63 |
| **HPYLSS1_00355** | Hypothetical protein | 7.82 |
| **HPYLSS1_01063** | Hypothetical protein | 7.58 |
| **HPYLSS1_00488** | Hypothetical protein | 5.65 |
| **HPYLSS1_01091** | Hypothetical protein | 5.34 |
| **HPYLSS1_00197** | Hypothetical protein | 4.4 |
| **HPYLSS1_00109** | Hypothetical protein | 4.25 |
| **HPYLSS1_00933** | Hypothetical protein | 3.91 |
| **HPYLSS1_01183** | Hypothetical protein | 3.37 |
| **HPYLSS1_00583** | Hypothetical protein | 3.07 |
| **HPYLSS1_00404** | Hypothetical protein | 2.85 |
| **HPYLSS1_01474** | Hypothetical protein | 2.77 |
| **HPYLSS1_00984** | Hypothetical protein | 2.56 |
| **HPYLSS1_01009** | Hypothetical protein | 2.26 |
| **HPYLSS1_01271** | Hypothetical protein | 2.05 |
| **HPYLSS1_00558** | Hypothetical protein | 2.04 |
| **HPYLSS1_01019** | Hypothetical protein | 2.03 |
| **HPYLSS1_00777** | Hypothetical protein | 2.01 |
| **HPYLSS1_00529** | Hypothetical protein | 2 |

**Table 3.** Down-regulated genes in *H. pylori* SS1 biofilm (cutoff ratio ≤ -1 log2 fold change and p-value <0.05) using RNA-seq analysis, grouped by functional role categories.

| Locus Name | Putative identification | Fold Change |
| --- | --- | --- |
| **Cell envelope** |  |  |
| *murB* (HPYLSS1_01344) | UDP-N-acetylenolpyruvoylglucosamine reductase | -2.52 |
| *fliM* (HPYLSS1_00401) | Flagellar motor switch protein FliM | -2.25 |
| *fliI* (HPYLSS1_01346) | Flagellum-specific ATP synthase | -2.16 |
| *yohD-1* (HPYLSS1_00775) | Inner membrane protein YohD | -2.14 |
| **Cellular process** |  |  |
| *mreB* (HPYLSS1_01316) | Rod shape-determining protein MreB | -2.95 |
| *urea* (HPYLSS1_00068) | Urease subunit alpha | -2.28 |
| *groEL* (HPYLSS1_00013) | 60 kDa chaperonin | -2.17 |
| *hcpC* (HPYLSS1_01039) | Cysteine rich protein HcpC | -3.78 |
| *cmmA* (HPYLSS1_01486) | Polymer-forming cytoskeletal family protein | -2.58 |
| *typA* (HPYLSS1_00442) | GTP-binding protein | -2.02 |
| **Regulatory functions** |  |  |
| *hslV* (HPYLSS1_00733) | ATP-dependent protease subunit | -3.33 |
| HPYLSS1_00758 | Putative TrmH family tRNA/rRNA | -2.24 |
| **Translation** |  |  |
| *rplR* (HPYLSS1_01253) | 50S ribosomal protein L18 | -3.09 |
| *rpsE* (HPYLSS1_01252) | 30S ribosomal protein S5 | -3.06 |
| *rpsG* (HPYLSS1_01140) | 30S ribosomal protein S7 | -2.52 |
| *rpsC* (HPYLSS1_01262) | 30S ribosomal protein S3 | -2.49 |
| *rpsK* (HPYLSS1_01246) | 30S ribosomal protein S11 | -2.44 |
| *rplW* (HPYLSS1_01266) | 50S ribosomal protein L23 | -2.29 |
| *rplN* (HPYLSS1_01258) | 50S ribosomal protein L14 | -2.29 |
| *rplD* (HPYLSS1_01267) | 50S ribosomal protein L4 | -2.23 |
| *rplB* (HPYLSS1_01265) | 50S ribosomal protein L2 | -2.22 |
| *rplF* (HPYLSS1_01254) | 50S ribosomal protein L6 | -2.21 |
| *rpmF* (HPYLSS1_00189) | 50S ribosomal protein L32 | -2.14 |
| *rpsD* (HPYLSS1_01245) | 30S ribosomal protein S4 | -2.12 |
| *rplS* (HPYLSS1_01093) | 50S ribosomal protein L19 | -2.09 |
| *rplE* (HPYLSS1_01256) | 50S ribosomal protein L5 | -2.07 |
| *rplV* (HPYLSS1_01263) | 50S ribosomal protein L22 | -2.02 |
| *rpmG* (HPYLSS1_01151) | 50S ribosomal protein L33 | -2.02 |
| *fusA* (HPYLSS1_01139) | Elongation factor G | -2.23 |
| *tufa* (HPYLSS1_01152) | Elongation factor Tu | -2.14 |
| *yigZ* (HPYLSS1_01411) | Elongation factor | -2.05 |
| **Amino acid biosynthesis** |  |  |
| *trpB* (HPYLSS1_01240) | Tryptophan synthase beta chain | -2.68 |
| **Fatty acid and phospholipid metabolism** |  |  |
| *acpP_2* (HPYLSS1_00944) | Acyl carrier protein | -2.96 |
| *plsX* (HPYLSS1_00190) | Phosphate acyltransferase | -2.71 |
| *scoB* (HPYLSS1_00895) | 3-Oxoacid CoA-transferase, subunit B | -2.11 |
| **Biosynthetic of cofactors, prosthetic groups and carriers** |  |  |
| *birA* (HPYLSS1_01084) | Bifunctional ligase/repressor BirA | -3.29 |
| *ribH* (HPYLSS1_00002) | 6,7-dimethyl-8-ribityllumazine synthase | -2.47 |
| *folK* (HPYLSS1_00396) | 2-amino-4-hydroxy-6-hydroxymethyldihydropteridine pyrophosphokinase | -2.13 |
| *ggt* (HPYLSS1_01061) | Gamma-glutamyl transpeptidase | -2.06 |
| **DNA restriction, modification, recombination, and repair** |  |  |
| HPYLSS1_00145 | Recombinase A | -2.46 |
| **Energy metabolism** |  |  |
| *atpC* (HPYLSS1_01075) | ATP synthase epsilon chain | -2.78 |
| *atpE* (HPYLSS1_01164) | ATP synthase subunit c | -2.48 |
| *nifU* (HPYLSS1_00210) | NifU-like protein | -2.15 |
| *adhA* (HPYLSS1_00) | Alcohol dehydrogenase | -2.12 |
| *mdaB* (HPYLSS1_00836) | Modulator of drug activity B | -2.05 |
| **Purine, pyrimidine, nucleosides and nucleotide** |  |  |
| *pyrD_2* (HPYLSS1_01468) | Dihydroorotate dehydrogenase B (NAD(+)), catalytic subunit | -2.13 |
| **Hypothetical proteins** |  |  |
| **HPYLSS1_00188** | Hypothetical protein | -3.92 |
| **HPYLSS1_00885** | Hypothetical protein | -3.15 |
| **HPYLSS1_00325** | Hypothetical protein/Putative beta-lactamase | -2.95 |
| **HPYLSS1_00036** | Hypothetical protein/Putative Nucleoid-associated protein | -2.79 |
| **HPYLSS1_01458** | Hypothetical protein | -2.79 |
| **HPYLSS1_01225** | Hypothetical protein | -2.71 |
| **HPYLSS1_00259** | Hypothetical protein | -2.66 |
| **HPYLSS1_01321** | Hypothetical protein | -2.44 |
| **HPYLSS1_01060** | Hypothetical protein | -2.32 |
| **HPYLSS1_00657** | Hypothetical protein | -2.27 |
| **HPYLSS1_01143** | Hypothetical protein | -2.21 |
| **HPYLSS1_00296** | Hypothetical protein/Putative F0F1-ATPase subunit | -2.2 |
| **HPYLSS1_00945** | Hypothetical protein | -2.16 |
| **HPYLSS1_00057** | Hypothetical protein | -2.03 |
| **HPYLSS1_00569** | Hypothetical protein | -2.07 |

**Fig 4.**
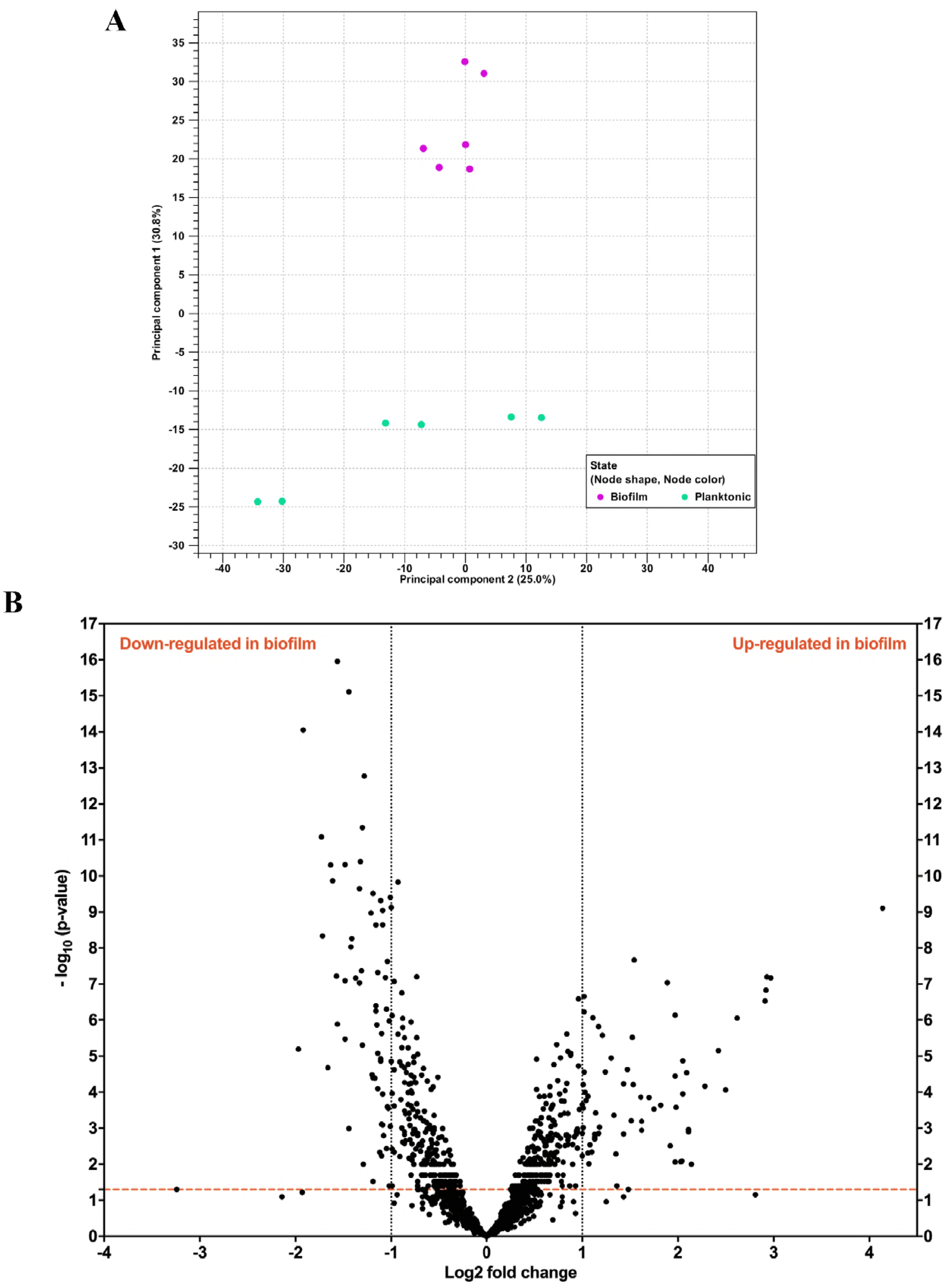
Biofilm-grown cells and planktonic cells show distinct transcriptional profiles. **(A)** principal component analysis (PCA) of gene expression obtained by RNA-seq between biofilm (n =3) and planktonic (n =3) populations. **(B)** Volcano plot of gene expression data. The y-axis is the negative log10 of P-values (a higher value indicates greater significance) and the x-axis is log2 fold change in difference in abundance between two population (positive values represent the up-regulated genes in biofilm and negative values represent down-regulated genes). The dashed red line shows where *P* =0.01, with points above the line having *P* < 0.01 and points below the line having *P* > 0.01.

**Fig 5.**
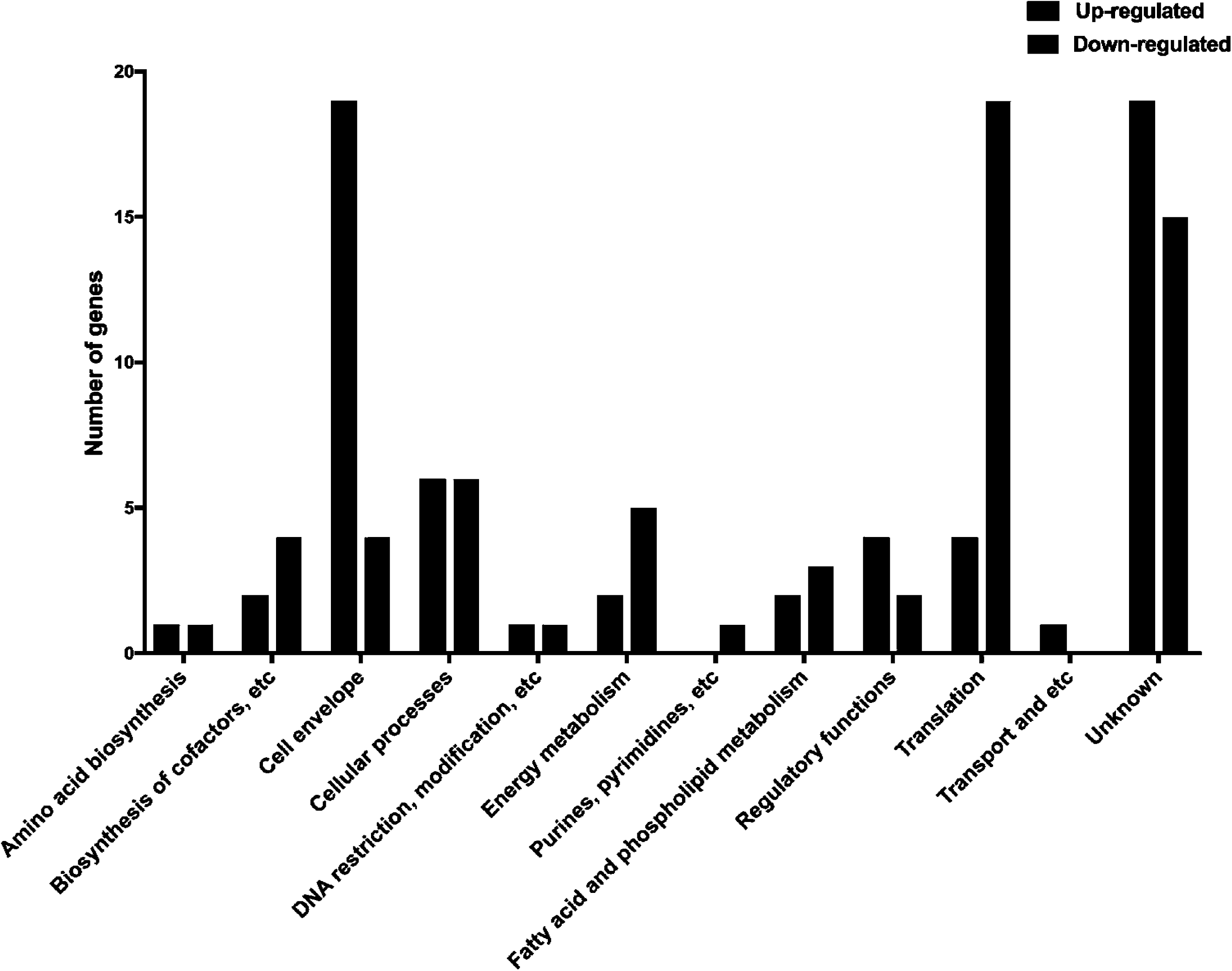
Functional classification of genes differentially expression in *H. pylori* SS1 biofilm. Black and grey bars represent up-regulated and down-regulated genes, respectively that were significantly differentially expressed (*p*< 0.01 and log2-fold change >1 or <-1) between *H. pylori* biofilm and planktonic populations

**Fig 6.**
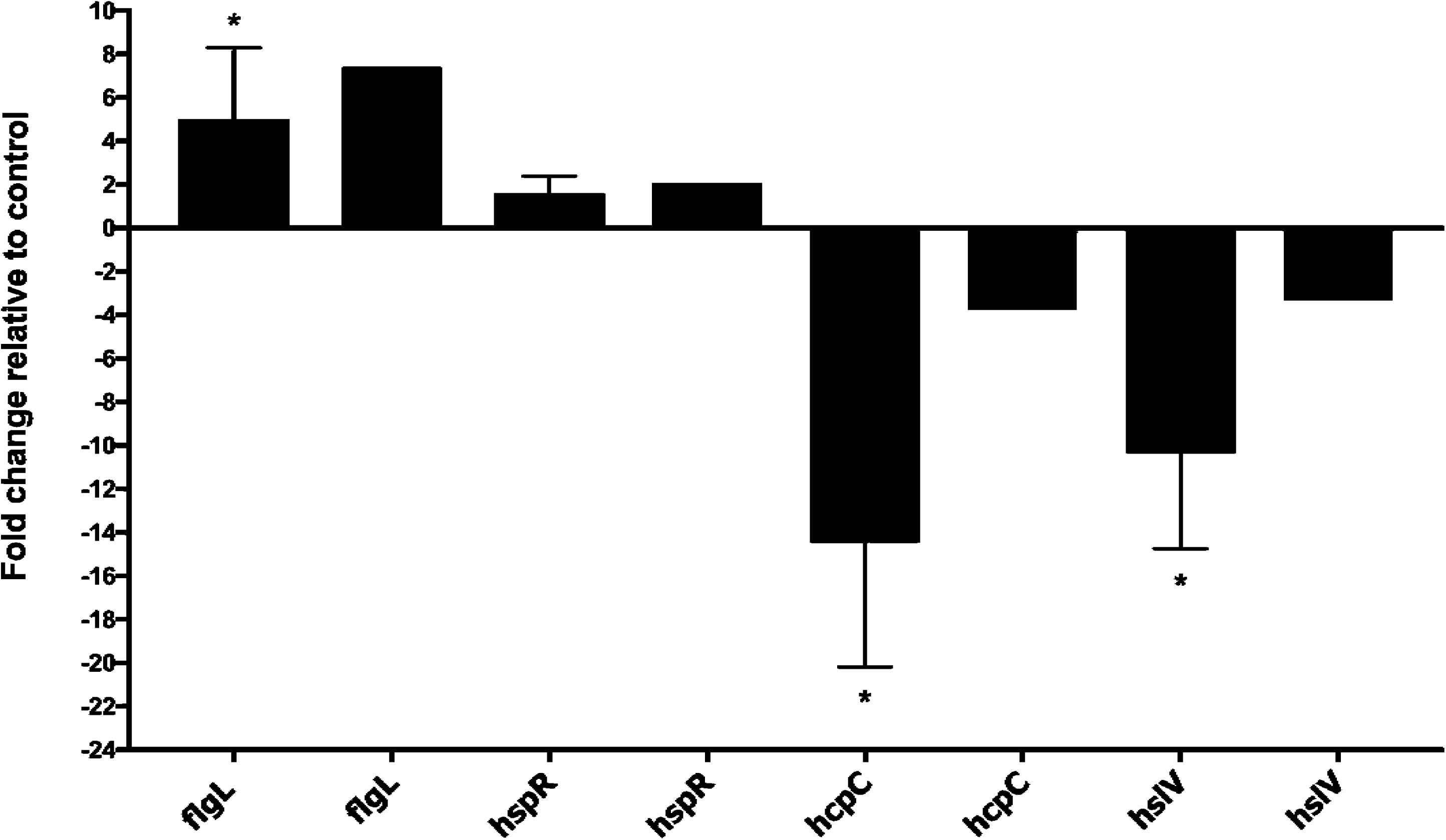
qPCR validation of the transcription of selected differentially expressed genes. The data indicate the fold-change expression of genes in *H. pylori* biofilm cells compared to planktonic cells. Fold-change in gene expressions were calculated after normalization of each gene with the constitutively expressed gene control *gapB*. Bars represent the mean and the error bars the standard error of the mean, Black and grey bars represent qPCR and RNA-seq results, respectively. Statistical analyses were performed using 2^-ΔΔ^*CT* values, and all results with an asterisk were statistically significant (*P < 0.01).*

We found evidence that biofilms cells experience a stressful environment. Indeed, genes coding for several stress response-related genes such as *hrcA, hspR, crdR, recR* and *pgdA* were up-regulated in biofilm cells (Table 2). The *hspR* and *hrcA* genes code for transcriptional repressor proteins belonging to the heat shock protein family, and were both up-regulated in biofilm cells. The *crdR* gene, which encodes a copper-related transcriptional response regulator was also up-regulated in *H. pylori* biofilm cells. Several transcripts encoding for oxidative stress resistance were similarly up-regulated in biofilm cells. These included *recR*, a gene encoding for a DNA recombination protein, as well as *pgdA* which encodes for a peptidoglycan deacetylase. These have both been previously associated with oxidative stress in *H. pylori* (33).

We found that the ATP-dependent protease HslV gene was among the most down-regulated genes in *H. pylori* biofilm (Table 3). Although this protein has not yet been studied in the context of *H. pylori* biofilms, the orthologous *E. coli* HslV protease has been previously associated with biofilm dispersal (34).

Our data suggest that biofilm cells may be less virulent in some ways, but more in others. Transcripts coding for some *H. pylori* virulence, colonization or immunogenic factors were low in biofilm cells, including the UreA subunit of urease, the GroEL chaperone, and the HcpC cysteine rich protein. These have each been shown to play roles in colonization or promoting inflammatory gene expression (35-37). On the other hand, only three genes encoded within the cytotoxin-associated gene pathogenicity island (*cag*PAI) (38,39) *cagL, cagW* and *cagE* were significantly highly expressed in biofilm cells of *H. pylori*. These genes are in separate operons (38), and encode for cag pathogenicity island protein CagL/Cag18, an integrin binding protein at the cag pilus tip, cag pathogenicity island protein CagW/Cag10 and type IV secretion system protein CagE/virB4, both part of the inner membrane protein transfer complex (39).

Many genes related to the cell envelope were up-regulated in biofilm cells (Fig. 5). Indeed, genes coding for proteins involved in lipopolysaccharide synthesis such as *lpxB*, which encodes a lipid-A disaccharide synthase, and *lptB*, which encodes a lipopolysaccharide export system ATPase, were up-regulated in biofilm cells (Table 2). Numerous transcripts encoding cytoplasmic and outer membrane proteins were also elevated in biofilm cells (i.e. *homC, homD* and HPYLSS1_00450) (Table 2).

Interestingly, the majority of the upregulated cell envelope genes in biofilm cells encoded for flagellar structure and biosynthesis proteins such as *flgL, flgK, fliD* and *flgE,* which encode for flagellar hook-associated proteins (Table 2). Two known or putative flagellin genes, *flaB* and *flaG* were also upregulated in the biofilms (Table 2). These data suggested the intriguing idea that flagella might play a role in the *H. pylori* biofilm.

### Flagella are present and play a structural role in H. pylori biofilms

The transcriptomic data above suggested that flagellar components are upregulated in the biofilm cells, so we used SEM to gain insights into the biofilm architecture of *H. pylori*. This analysis demonstrated three-dimensional structures composed of bacterial cells adherent to one another and to the surface (Fig. 7A). Biofilms contained mainly coccoid cells along with some rod-shaped cells (Fig. 7A), as described previously for *H. pylori* biofilms (20-23). When compared to planktonic populations, the proportion of both morphologies were similar at ∼ 80% coccoid cells (data not shown).

Interestingly, extensive networks of bundles of filaments were visible in the biofilms. In some cases, these appeared to be connected to the bacterial pole, as would be expected for flagella (Fig. 7A, arrowheads). We measured the dimensions of the filaments to see if they were consistent in size with flagella. The width and the length measured at 20 to 30nm and 3 to 4μm, respectively, and were in agreement with those reported previously for *H. pylori* flagella (40). This data, especially when combined with transcriptomics, suggested these structures could be flagella. We therefore analyzed a mutant strain that lacks a key component of the flagellar basal body, FliM, and is aflagellated (41). SEM analysis of the Δ*fliM* mutant showed a complete loss of flagella (Fig. 7A). This mutant displayed significantly less biofilm biomass (Fig. 7B), but we were able to find few microcolonies. Within these microcolonies, the filaments were completely lacking (Fig. 7A). These results suggest that these filaments are flagella, and furthermore that flagella and/or motility are important for biofilm formation.

**Fig 7.**
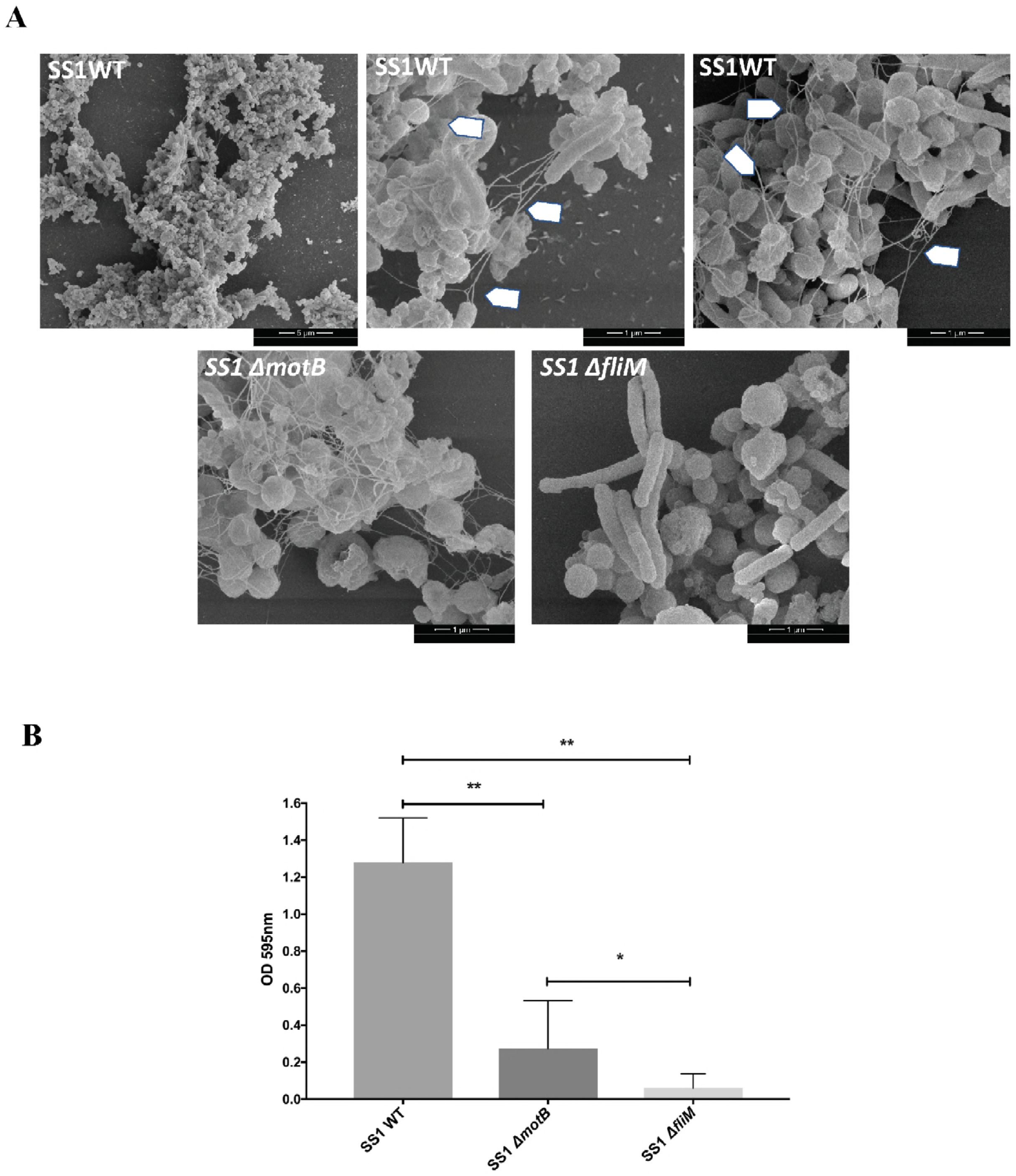
Flagella play integral roles in *H. pylori* biofilms. **(A)** Scanning electron microscope (SEM) images of biofilms formed by *H. pylori* wild-type SS1 (SS1 WT), isogenic non-motile but flagellated mutant Δ*motB* (SS1 Δ*motB*), and isogenic aflagellated mutant Δ*fliM* (SS1 Δ*fliM*). Arrows flagella. **(B)** Quantification of biofilm formation by *H. pylori* SS1 WT, Δ*motB* and Δ*fliM*. Strains were grown in BB2 media for three days, followed by biofilm evaluation using the crystal violet assay. Experiments were performed three times independently with 6-9 technical replicates for each. Statistical analysis was performed using ANOVA (**, *P* < 0.01 and *, *P* < 0.05).

To further dissect the roles of motility and flagella in biofilm formation, we analyzed biofilm formation in a non-motile but flagellated strain created by disruption of the motor protein MotB. As reported previously, this mutant still expresses flagella (Fig. 7A). Biofilm formation, however, was severely impaired compared to the wild-type strain, which suggest that a lack of motility might contribute to the biofilm defect. However, Δ*motB* produced significantly more biofilm than Δ*fliM* mutant suggesting that the flagella structure even in absence of motility also contributes to biofilm formation in *H. pylori*.

To examine whether other strains of *H. pylori* similarly use flagella in biofilms, we imaged the biofilm of *H. pylori* strain G27 and similar flagellar mutants as used above. Wild-type *H. pylori* G27 biofilm cells also contained filaments consistent with flagella (Fig. 8A). As with strain SS1, mutants lacking flagella (*flgS or fliA*) formed very weak biofilms, while strains that had flagella but no motility (*motB*) retained partial biofilm formation (Fig. 8B).

**Fig 8.**
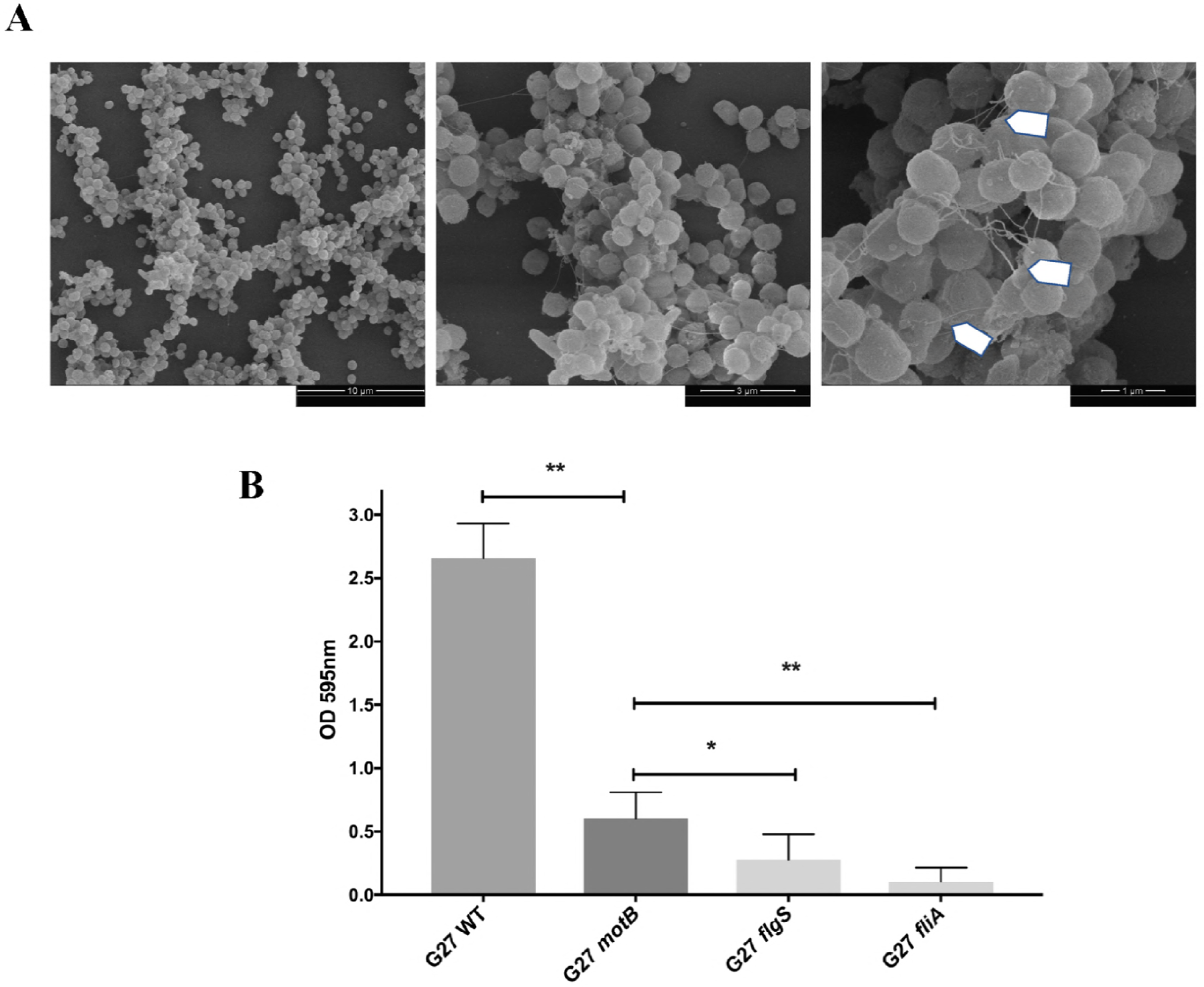
*H. pylori* G27 biofilm contain structurally important flagella. **(A)** Scanning electron microscope (SEM) images of wild type G27 *H. pylori* biofilms. Arrows indicate flagella. **(B)** Quantification of biofilm formation by *H. pylori* G27 wild type (WT), non-motile flagellated *motB*, non-motile mutant *fliA* that is reported to have either truncated flagella or no flagella, and the aflagellated and non-motile mutant *flgS*. Biofilms were evaluated using the crystal violet assay. Experiments were performed 2 times independently with at least 6 technical replicates for each. Statistical analysis was performed using ANOVA (**, *P* < 0.01 and *, *P* < 0.05).

Taken together, these data suggest that flagella are produced by *H. pylori* when in a biofilm, and appear to play roles in addition to simple motility, promoting biofilm integrity by holding cells together and to the surface,

## DISCUSSION

In this report, we present the first transcriptomics characterization of the *H. pylori* biofilm. This study demonstrated clearly distinct expression profiles between planktonic and biofilm cells. The biofilm cells were characterized by low metabolic activity and triggering of several stress responses. Among the upregulated genes in the biofilm cells, we found several genes associated with cell membrane proteins, outer membrane proteins, stress response, and surprisingly, genes related to the flagellar apparatus. SEM analysis confirmed that flagella are present in a mature *H. pylori* biofilm, and appear to play a role in maintaining solid biofilm structures. This result was somewhat surprising, as typically flagella are proposed to be turned off during the sessile biofilm growth mode (42-44). Recent work however, discussed below, has suggested that flagella in *E. coli* biofilms may play a structural role. Our studies with *H. pylori* thus build on an emerging theme that flagella are not always turned off in mature biofilms, and indeed may play important functions in biofilm structure

To gain insights into the mechanisms behind the biofilm formation in *H. pylori,* we used RNA sequencing and carried out a comparative transcriptomic analysis between biofilm cells and those in the planktonic state. Using this approach, we observed that 8.18% of genes were significantly differentially expressed between biofilm and planktonic cells, similar to that reported in other bacterial systems (43,45,46). In our experimental design, we compared a static biofilm mode of growth, where attached cells adhered to the bottom of the wells, with planktonic non-attached cells in the same wells. This approach was used to maintain the same growth conditions as much as possible between biofilm and planktonic samples, and likely contributed to the relatively small number of differentially expressed genes. However, since biofilm formation is a dynamic process with frequent switching between planktonic to biofilm modes occurring frequently, we likely have some contamination between the biofilm and planktonic populations. Therefore, our method may have missed some genes that are expressed in either population.

One of the findings from our transcriptomic analysis was that several flagellar protein transcripts were significantly elevated in the biofilm. Notably, these were not for the entire flagellum, but instead specific genes encoding the rod, hook, and filament. Specifically, we saw biofilm-cell overexpression of genes encoding for the FlgB rod protein, the FlgE flagellar hook protein, the FlgK and FlgL hook-filament junction proteins, the FliK hook length control protein and two flagellins (FlaB and the putative flagellin encoded by FlaG). Notably absent was the gene for the major flagellin FlaA, and genes for the motor and stator. We also saw the up-regulation of *flgM* which encodes an anti-sigma factor that interacts with flagellar sigma factor FliA, and therefore would be expected to decreased expression of *flaA* (47).

Historically, flagella have been typically viewed as important only for initial biofilm attachment and later cell dispersion (44,48). In fact, it has often been suggested that genes encoding for flagella are turned off in mature biofilms (42-44). However, other reports have shown that some microbes express flagella during all stages of biofilm development and not only during the attachment and dispersion processes (49,50). In *E. coli*, several flagellar-biosynthesis genes were induced in mature biofilms, and around 20 flagellar genes were regulated throughout all stages of biofilm development and not simply turned off (49). *E. coli* flagella were proposed to have a structural role along with other matrix components (i.e. eDNA and extracellular proteins), acting to cement and hold cells together and to the surface (50,51). Our data furthermore showed that aflagellated mutants are poor biofilm formers, supporting that these filaments could play a structural role. Taken together, these findings suggest that flagella of *H. pylori* may play a structural role during biofilm formation to help bacteria attach to each other and to surfaces.

Interestingly, we found that the HspR and HrcA transcriptional repressor proteins are up-regulated in biofilm cells. These proteins had previously been shown to positively correlate with flagella expression (52), providing candidate regulatory proteins that function in biofilm cells. HspR and HrcA belong to the heat shock protein family, and have been shown to respond to heat shock temperature conditions although the nature of their “true” signal is not yet clear (52,53). A previous comparative transcriptomic analysis of wild-type *H. pylori* along with Δ*hspR,* Δ*hrcA*, and double mutants revealed a set of 14 genes that were negatively regulated and 29 genes that were positively regulated by these transcriptional regulators (52). The regulated genes include those for chaperones, urease enzyme activity, adhesion to epithelial cells and flagella. Interestingly, among the 29 positively regulated genes, nearly half (14) encoded for flagellar genes, including the *flgM, flaG, fliD, flgK, flgB, flgE* and *fliK* transcripts we identified here. Thus, our data suggest that biofilm conditions activate expression of HrcA and HspR, which in turn upregulate a subset of flagellar genes.

Experiments suggest that HrcA and HspR regulators do not directly regulate the flagellar genes (52). However, they do directly repress expression from several promoters including those upstream of the *groESL, hrcA-grpE-dnaK*, and *cbpA-hspR-hp1026* operons. These gene products encode the major chaperones of *H. pylori* (52-54). Heat shock conditions relieve the repression, and allow expression of these operons. Consistent with elevated expression of HspR and HrcA, we found the genes coding for the heat shock protein GroEL to be downregulated in biofilm. Our data suggest that some yet-to-be determined conditions occurring during biofilm formation trigger the expression of HspR and HcrA regulators.

Other genes associated with stress responses were also up-regulated in biofilm cells including the *pgdA* and *recR* genes. These genes encode for a peptidoglycan deacetylase and DNA recombination protein, respectively. RecR has been shown to be involved in repairing in DNA double strand breaks induced by oxidative stress (55) and the *recR* mutant was highly sensitive to DNA damaging agents, oxidative stress and had a reduced ability to colonize mouse stomachs (55). *pgdA* has been reported to be highly induced by oxidative stress (33,56). Up-regulation of oxidative stress genes has previously been reported in biofilm cells of other organisms including *E. coli* (57), *Pseudomonas aeruginosa* (42), *Neisseria gonorrhoeae* (58) and *Clostridium perfringens* (48).

As reported for other microorganisms, *H. pylori* biofilm cells have altered metabolism, typically thought to be associated with the restricted availability of nutrients (48,59). *H. pylori* biofilm cells were characterized by a downregulation of the expression of multiple genes involved in metabolism and translation including, *atpC, atpE, nifU* and several ribosomal protein genes. This low metabolism phenotype seems not be related simply to the presence of coccoid cells, but rather to the microenvironment generated during biofilm formation since the proportion of rods and coccoid forms did not differ between planktonic and biofilm populations.

*H. pylori* biofilm cells may also actively block the translational machinery, as suggested by the up-regulation of the gene encoding RsfS, a ribosomal silencing factor. This protein was previously described in *E. coli* and *Mycobacterium tuberculosis* to slow or block the translation machinery during stationary phase and/or nutrition-deficiency stress (60). It interacts with the 50S large ribosomal subunit, prevents its association with the 30 S ribosomal submit, and thus blocks formation of functional ribosomes (60). Whether it functions similarly in *H. pylori* remains to be determined.

These observations that biofilm cells may have decreased translation are relevant because at least two of the main antibiotics used to treat *H. pylori* infection, clarithromycin and tetracycline, inhibit the 50S and 30S ribosomal subunits, respectively. Thus, these antibiotics may have less impact on biofilm cells. In fact, recent *in vitro* studies have shown that clarithromycin is 4 to 16-fold less effective on *H. pylori* biofilm cells as compared to planktonic ones (26,61).

Taken together, our study has shown that *H. pylori* biofilm cells display a distinct transcriptomic profile compared to their planktonic counterparts. Lower metabolism and stress responses, likely associated to the microenvironment generated in the *H. pylori* biofilm, could be determinants of antimicrobial tolerance and involved in the persistence and survival of *H. pylori*. However, the up-regulated and down-regulated genes identified in this study are not specific for biofilm cells, and stress response genes have been previously observed in other conditions when both planktonic or biofilm cells were exposed to various stresses. Therefore, our data do not support the existence of a biofilm-specific genetic program. Additionally, our data show that flagella filaments are upregulated in biofilm cells and form an integral part of the biofilm matrix. Indeed, *H. pylori* without flagella form weak biofilms. These results thus contribute to correcting the idea that flagella are only involved during the first and last steps of biofilm formation, and instead support their importance throughout the biofilm process.

## MATERIALS AND METHODS

### Bacterial strain and growth conditions

*H. pylori* Sydney strain 1 (SS1) (62) and all other *H. pylori* strains used in this study are listed in Table.1. Strains were grown on Columbia Horse Blood Agar (CHBA), containing: 0.2%-β-cyclodextrin, 10μg/ml vancomycin, 5μg/ml of cefsulodin, 2.5 U/ml polymyxin B, 5μg/ml trimethoprim, and 8μg/ml amphotericin B (all chemicals from Thermo Fisher or Gold Biotech). Cultures were grown under micro-aerobic conditions (5% O_2_ and 10% CO_2_) at 37°C. For liquid culture and biofilm assay, *H. pylori* was grown in Brucella broth (Difco) containing 10% heat inactivated fetal bovine serum (FBS) (BB10; Gibco/BRL) with constant shaking under microaerobic conditions. For biofilm formation, several conditions were tested including Brucella broth containing different percentage of FBS (BB2, BB6 and BB10), and HAM’s F-12 (PAA Laboratories GmbH, Pasching, Austria) containing 10% or 2% of FBS (HAMS10 and HAMS2).

### Biofilm assays

Biofilm formation assays were carried as described previously, with slight modification (27). *H. pylori* SS1 was grown overnight in BB10 as above, diluted to and OD600 of 0.15 with fresh BB10, BB2, BB6 or HAMS media as desired, and then used to fill triplicate wells of a sterile 96-well polystyrene microtiter plate (Costar, 3596). Following static incubation of 1, 2, 3 or 5 days under micro aerobic conditions, culture medium was removed by aspiration and the plate was washed twice using PBS. The wells were then filled with 200μL of crystal violet (0.1 % wt/vol), and the plate was incubated for 2 min at room temperature. After removal of the crystal violet solution by aspiration, the plate was washed twice with PBS and dried for 20 min at room temperature. To visualize biofilms, 200 μL of ethanol (70% vol/vol) was added to the wells and the absorbance at 590 nm was measured.

### Biofilm dispersion assays

To evaluated the composition of SS1 biofilm matrix, we assessed the response of preformed biofilms to different enzymatic treatment. DNAse I and proteinase K (both from Sigma-Aldrich) were used to target, extracellular DNA and extracellular proteins, respectively. Biofilms were grown as described above and after three-days of growth, the old media were replaced by fresh media containing different concentrations of DNase I (380μg/ml to 95μg/ml) or proteinase K (200ug/ml to 50 ug/ml). The cells were then incubated for a further 24 hours. Control wells were exposed to media without enzyme. After treatments, the biofilm was stained with crystal violet as described above. Results are presented a percentage of the untreated control.

### Confocal laser scanning microscopy

Biofilms of *H. pylori* SS1 were prepared as described above using BB2, however, for confocal laser scanning microscopy (CLSM), μ-Slide 8-well glass bottom chamber slides (ibidi, Germany) were used instead of 96-well microtiter plates. Three day-old biofilms were stained with FilmTracer™ FM^®^1–43 (Invitrogen), BOBO-3 (Invitrogen), Filmtracer SYPRO Ruby biofilm matrix stain (Invitrogen), or FilmTracer LIVE/DEAD biofilm viability kit (Invitrogen) according to the manufacture’s instructions. Stained biofilms were visualized by CLSM with an LSM 5 Pascal laser-scanning microscope (Zeiss) and images were acquired using Imaris software (Bitplane). Biomass analysis of biofilm was carried using FM^®^1–43 stained z-stack images (0.1μm thickness) obtained by CLSM from randomly selected areas. The biomass of biofilms was determined using COMSTAT (63).

### RNA extraction and library construction

Biofilms of *H. pylori* SS1 were grown in 6-well plates (Costar) in BB2 as above. After 3 days of incubation, media containing non-attached planktonic bacteria (the planktonic fraction) was removed by pipetting, the cells were harvested by centrifugation and washed twice with PBS. The attached bacteria, representing the biofilm fraction, were washed twice with PBS to remove any remaining planktonic cells. Attached cells were scrapped off the plate using cell scraper. Both planktonic and biofilm fractions were subject to total RNA extraction using Trizol Max bacterial enhancement kit (Ambion, Life Technology, Carlsbad, CA, USA) as described by the manufacturer. RNA was further purified and concentrated using an RNAeasy Kit (Qiagen). rRNA was removed using the RiboZero magnetic kit (Illumina). Sequencing libraries were generated using NEBNext Ultra^TM^ Directional RNA library Prep Kit for Illumina (NEB, USA). cDNA library quality and amount was verified using Agilent Bioanalyzer 2100 system (Agilent technologies, CA, USA) and then sequenced using Illumina NextSeq Mid-Output (UC Davis Genome Center).

### Transcriptomic analysis

RNA-seq data were analyzed using CLC Genomics Workbench (Version 11.0, CLC Bio, Boston, MA, USA). After adapters were trimmed, forward and reverse sequenced reads generated for each growth state (biofilm vs planktonic; three biological replicates for each condition) were mapped against the SS1 reference genome (32) to quantify gene expression levels for each experimental condition. The expression value was measured in Reads per Kilobase Per Million Mapped Reads (RPKM). Genes were considered as differentially expressed when the log2 (fold change) was above 1 and the *P*-value was lower than 0.05.

### Quantitative PCR

To validate the RNA-seq data, we performed qPCR to quantify the transcription of four differentially expressed genes (two up-regulated genes and two down-regulated genes). The Fold change in gene expression was calculated after normalization of each gene with the constitutively expressed gene *gapB* (64). Primers used for this experiment are listed 5’-3’ below: gapB forward: GCCTCTTGCACGACTAACGC; gapB reverse: CTTTGCTCACGCCGGTGCTT. flgL forward: CAGGCAGCTCATGGATGCGA; flgL reverse: CGCTGTGCAAGGCGTTTTGA; hspR forward: TAGGCGTGCACCCTCAAACC; hspR reverse: CGCCCGCTAGATTAACCCCC; hcpC forward: GGGTTTTGTGCTTGGGTGCG; hcpC reverse: TTCCACCCCCTGCCCTTGAT; hslV forward: GATTTGCCGGAAGCACTGCG; hslV reverse: ATCATCGCTTCCAGTCGGCG

### Construction of H. pylori mutants

The SS1 Δ*fliM* mutant was created by natural transformation of SS1 wild type with plasmid pBS-fliM∷catmut (40) which replaces most of the fliM gene, corresponding to amino acids 1-105, with cat. The G27 *motB* mutant was created by natural transformation of G27 wild type with plasmid pKO114K and selection for kanamycin resistance. pKO114K was made as described for pKO114i (65), but instead of insertion of a aphA3-sacB allele, only an aphA3 allele was inserted. This allele inserts the aphA3 at the position corresponding to amino acid 113 of 257.

### Scanning electron microscopy

*H. pylori* biofilms were grown on glass coverslips (12mm, Chemgalss, life Sciences, Vineland, NJ) by dispersing 4 mL of a culture diluted to OD 0.15 in BB2 into wells of a 6-well plate (Costar). The plate was incubated as described above. After three days of growth, biofilms formed on the surface of the coverslips and planktonic cells were washed twice with PBS and fixed with 2.5% glutaraldehyde (wt/vol) for 1 hour at room temperature. Samples were then dehydrated with graded ethanol series, critically point dried, sputtered with ∼20 nm of gold (Hammer IV, Technics Inc, Anaheim, CA) and imaged in an FEI Quanta 3D Dualbeam SEM operating at 5 kV and 6.7 pA.

### Statistical analysis

Biofilms data were analyzed with GraphPad Prism (version 7.0) software (GraphPad Inc., San Diego, CA) using one-way analysis of variance (ANOVA) followed by Dunnett’s multiple-comparison test.

## ACKNOWLEDGEMENTS

We thank Dr. Fitnat Yildiz (U.C. Santa Cruz) and Dr. Davide Roncarati (University of Bologna, Italy) for their helpful suggestions and comments on the study and the manuscript. We thank Aaron Clarke (U.C. Santa Cruz) for his assistance with portions of the SEM experiments. We thank Dr. Ben Abrams (U.C. Santa Cruz) for CLSM assistance. We also acknowledge Dr. Tom Yuzvinsky (U.C. Santa Cruz) for assistance with sample preparation and electron microscopy and the W.M. Keck Center for Nanoscale Optofluidics for use of the FEI Quanta 3D Dualbeam microscope. We also thank Dr. David Bernick and METX department (U.C. Santa Cruz) for holding and financially supporting the RNAseq workshop. The work described here was supported by National Institutes of Health National Institute of Allergy and Infectious Disease (NIAID) grant RO1AI116946 (to K.M.O.). The funders had no role in study design, data collection and interpretation, or the decision to submit the work for publication.

---

**Supplementary Movie 1. Three-dimensional (3D) view of *H. pylori* biofilm grown for 3 days.** Bacteria were stained with FilmTracer™ FM^®^1–43 and observed by CLSM.

